# Aneuploid embryos as a proposal for improving Artificial Intelligence performance

**DOI:** 10.1101/2022.11.16.516774

**Authors:** Enric Güell Penas, Marina Esquerrà Parés, Andreu Vives Perelló, Mikaela Mladenova Koleva

**Affiliations:** CONSULTFIV, Pg. Estació 18 5-5, 43800 Valls, Spain; Institut Conceptum, Pg. Sunyer 49, 43202 Reus, Spain

**Keywords:** Artificial Intelligence, Machine Learning, Morphokinetics, Morphodynamics, Aneuploidy

## Abstract

**RESEARCH QUESTION:** Could we improve the performance of Machine Learning algorithms by using aneuploid embryos instead of non-implanted embryos as the contrary reference to Live-Birth embryos?

**DESIGN:** A single-center retrospective analysis of 343 embryos through 3 ML algorithms, based on manually annotated morphokinetics from Day 1 to Day 3. Two datasets were built including the same Live-Birth embryos (117). Dataset A included 123 non-implanted embryos, while Dataset B included 103 aneuploid embryos. V-Fold Cross-Validation was performed for each dataset and algorithm and the Area Under the Curve (AUC) was registered.

**RESULTS:** AUC for Dataset A did not reach 0.6 for any of the algorithms; while AUC values for “Dataset B” surpassed 0.7. According to this, different morphokinetic patterns were detected by Machine Learning algorithms.

**CONCLUSIONS:** Algorithms’ minor performance with non-implanted embryos may be due to an increased Label Noise effect, suggesting that including aneuploid embryos could be more appropriate when building predictive algorithms for embryo viability. Machine Learning algorithms results were improved when aneuploid embryos were taken into consideration.

## INTRODUCTION

Artificial Intelligence (AI) and Machine Learning (ML) in the *in vitro* fertilization (IVF) lab is increasing the presence everyday as more utilities are being found to have a promising predictive power (Simopolou et al., 2018; Wang et al., 2019; Zaninovic et al., 2020; Riegler et al., 2021) although there is still room for improvement (Curchoe et al., 2020; Rosenwaks et al., 2020; Swain et al., 2020; Dimitriadis et al., 2022). When working with supervised models (Morales et al., 2008; Fernandez et al., 2020) embryologists are responsible for reporting to the machine the level at which belongs every embryo. Unproperly labelling could lead to distorted predictions and decreasing in the AI performance (Zhu & Wu, 2004; Frénay & Verleysen, 2014), as the machine would be looking for differences between embryos that, in fact, could belong to the same group.

Euploidy is not an absolute guarantee for the embryo to have high developmental potential (Irani et al., 2016; Murugappan et al., 2020) and mosaicism cannot be discarded (Popovic et al., 2018; Griffin & Ogur, 2018). Actually, the only mislabelling-free statement is that Live-Birth category is composed by optimal potential embryos containing at least a minimum euploid load allowing them to continue their proper development. Non-implanted embryos, which are used in traditional Live-Birth prediction (Vermilyea et al., 2020), could involve a certain number of potentially evolving embryos with a negative result due to factors unrelated to the embryo (Weimar et al., 2013; Atwood & Adakkadath, 2016; Milewski et al., 2017a; Afnan et al., 2021). Finally, aneuploidy is a proof of genetic dysfunction despite some aneuploid embryos could still be viable due to undetected mosaicism or non-concordance diagnosis (Victor et al., 2019).

Therefore, the aim of this study was to analyse if the performance of Machine Learning algorithms could improve by using aneuploid embryos instead of non-implanted embryos as the contrary reference to Live-Birth embryos.

## MATERIALS AND METHODS

### Study Population

This project was a retrospective observational study of 343 embryos carried out at Institut Conceptum IVF lab in Reus (Tarragona, Spain) with the technical and conceptual support of ART Consultancy CONSULTFIV from Valls (Tarragona, Spain). This study and its protocol were approved by the Ethics Committee of Hospital Clinic de Barcelona (reference HCB/2021/1006).

### Experimental Design

The study consisted on comparing the performance of good morphology blastocysts (Zhan et al., 2020) labelled as Live-Birth, Non-implanted and Aneuploid through three different Machine Learning algorithms. Study groups of this work are schematically outlined in Figure 1. All the embryos were cultured inside a conventional incubator (Labotect C-200, Labotect Labor-Technik-Göttingen GmbH, Germany), (37°C, 6% CO2), equipped with time-lapse technology (TLT) Primo Vision (Vitrolife, Gothenburg, Sweden). The inclusion criteria for Live-Birth prediction was checking the newborns healthy Live-Birth status. Aneuploid diagnosis was obtained by Preimplantation Genetic Testing for Aneuploidies (PGT-A) of a single blastomere on Day 3. Two different datasets were built sharing the same Live-Birth embryos (n = 117). Dataset “A” included non-implanted embryos (n = 123) while Dataset “B” comprised aneuploid embryos (n = 103). Baseline characteristics of the study groups included in the datasets are showed at Table 1.

**Figure 1.**
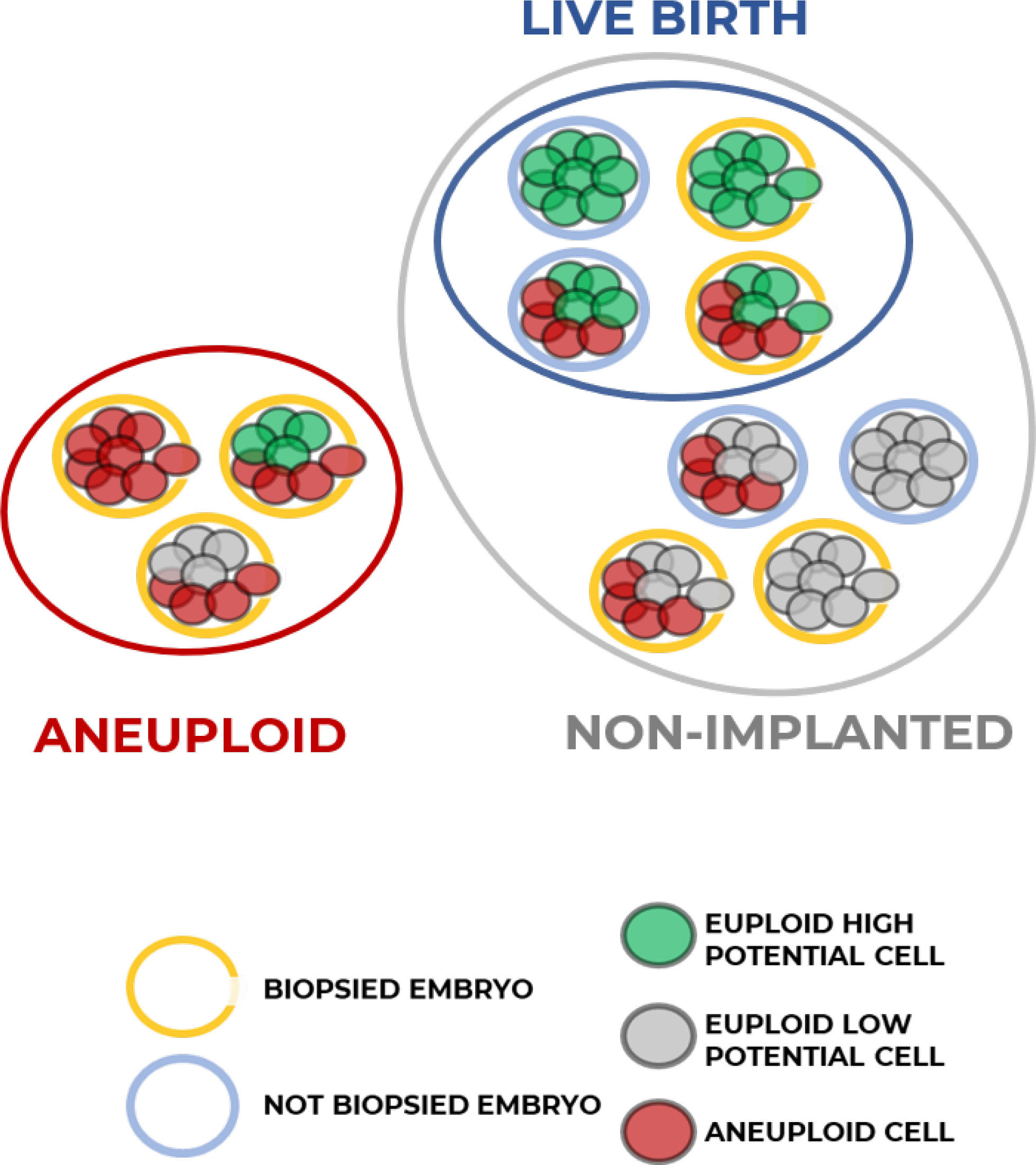
Studied groups of the outcome variable Embryo Result (Live-Birth, Non-implanted and Aneuploid).

### Morphokinetic analysis

Manually annotated morphokinetic and morphodynamic analysis was performed on Primo Vision Analyzer software. Time of ICSI was taken into consideration for every oocyte (Ciray et al., 2014) and morphokinetic time points were relativized for each embryo at its own t0 microinjection time (Güell et al., 2019) by CONSULTFIV’s platform named “TLReader”. Morphokinetic parameter (MCP) annotation was also based on 2014 Ciray’s publication: polar body exclusion (tPB2), pronuclear fading (tPNf), embryonic cell cleavages (t2-t8) as shown in Figure 2. Nevertheless, there were also included novel biomarkers considering the fading of the nucleus at the 2-cell stage (tNf Cc2) of the first cell cleaving to t3 and at the 4-cell stage (tNf Cc3) of that first cell which cleaved to t3 on the previous cell cycle and which is going to cleave at t5. Morphodynamic phenomena (MDp) were marked with the type of MDp and the cell cycle (CC) at which occurred, and t-times adjustments were performed in order to maintain the traceability of each cell lineage. These adjustments were customised depending on the type and cell cycle of MDp. An example in a diagram for the tripolar mitosis at CC2 adjustment is presented in Video 1.

**Figure 2.**
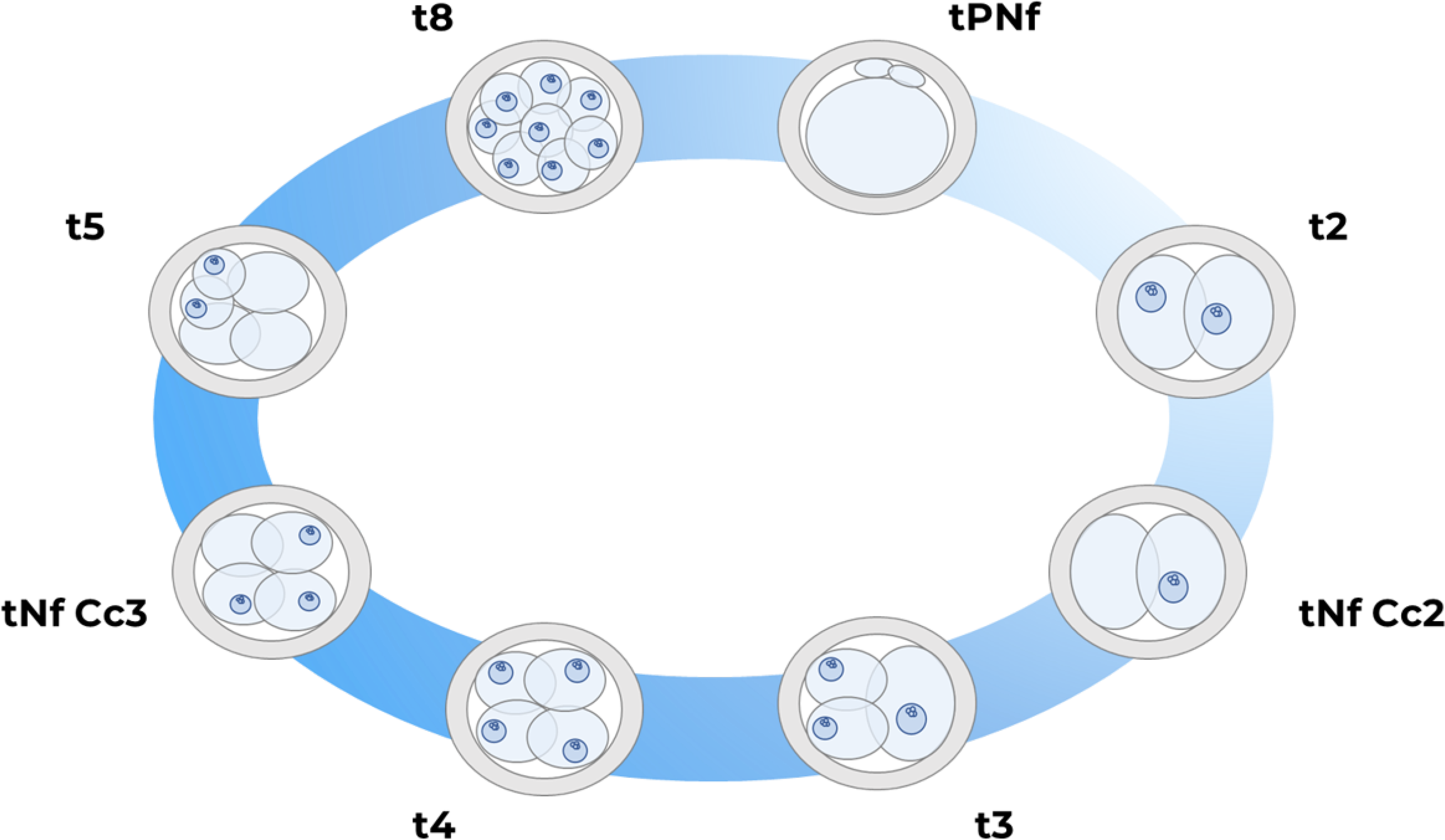
Morphokinetic parameters annotated in the present project.

### Machine Learning algorithms

Three machine learning algorithms (Rudin et al., 2021) were trained and v-fold cross-validated by 80% train set, 20% validation set (Kuhn & Johnson, 2013): an eXtreme Gradient Boosting (XGB), a k-Nearest Neighbor (kNN) and a Random Forest (rF). After feature extraction, we decided to build each algorithm with 4 biomarkers related to the cell lineage of cell “A” (Figure 3): tNf Cc2, t3, tNf Cc3 and t5. The prediction power for each dataset model was measured using the area under the receiver operating characteristic (ROC) curve (AUC), ranging from 0.5 (completely random choices) to 1.0 (perfect discrimination) and its confusion matrix’s metrics.

**Figure 3.**
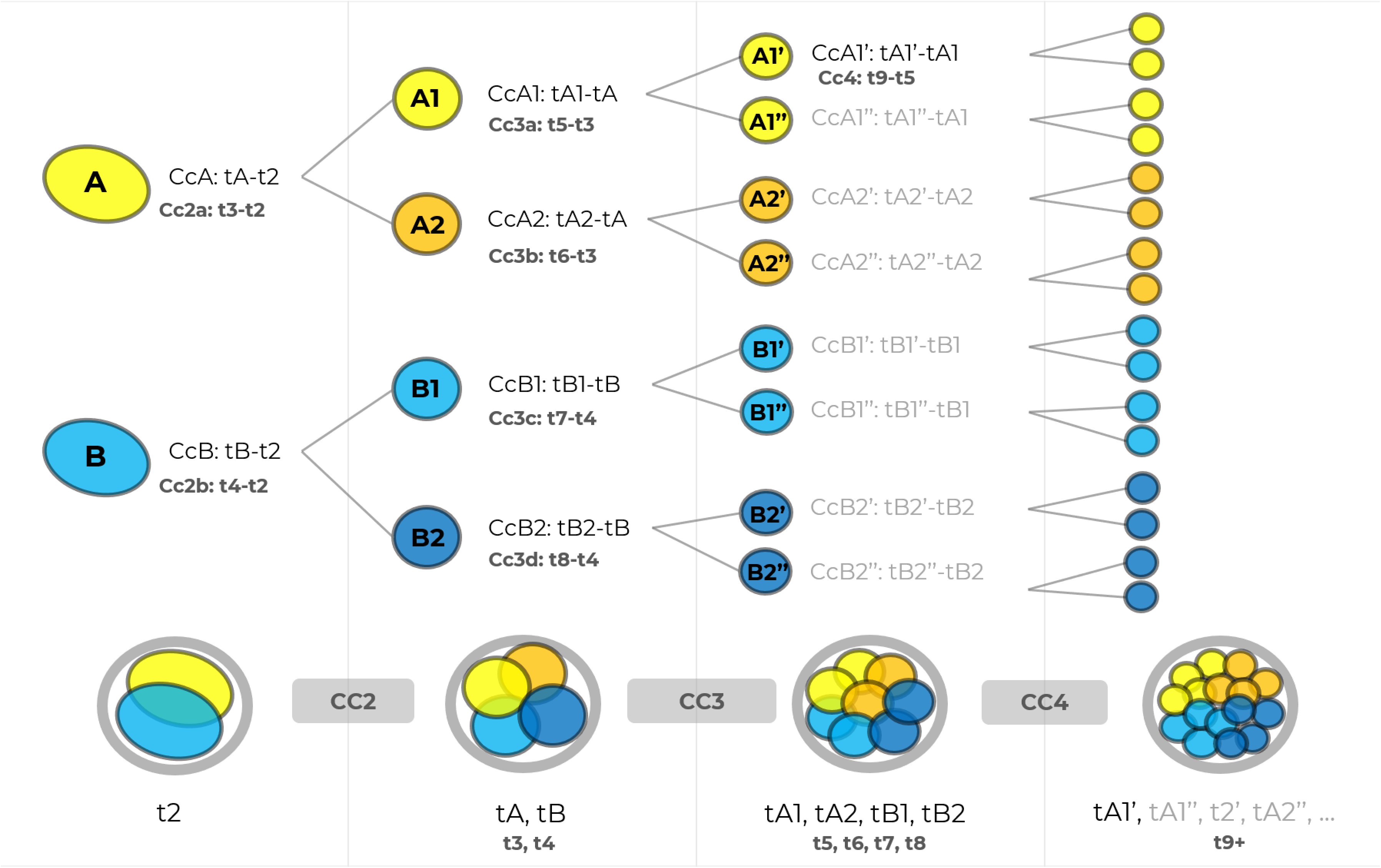
Schematic diagram about naming cells depending on the cell lineage.

## RESULTS

Regarding the results, all metrics for each Machine Learning algorithm analysed were higher in the Dataset including aneuploid embryos (Dataset B) rather than in the Dataset including non-implanted embryos (Dataset A). The AUC for Dataset “A” did not reach the value of 0.6 for any of the three algorithms while the AUC for dataset “B” surpassed the value of 0.7 for each one of them (Figure 4).

**Figure 4.**
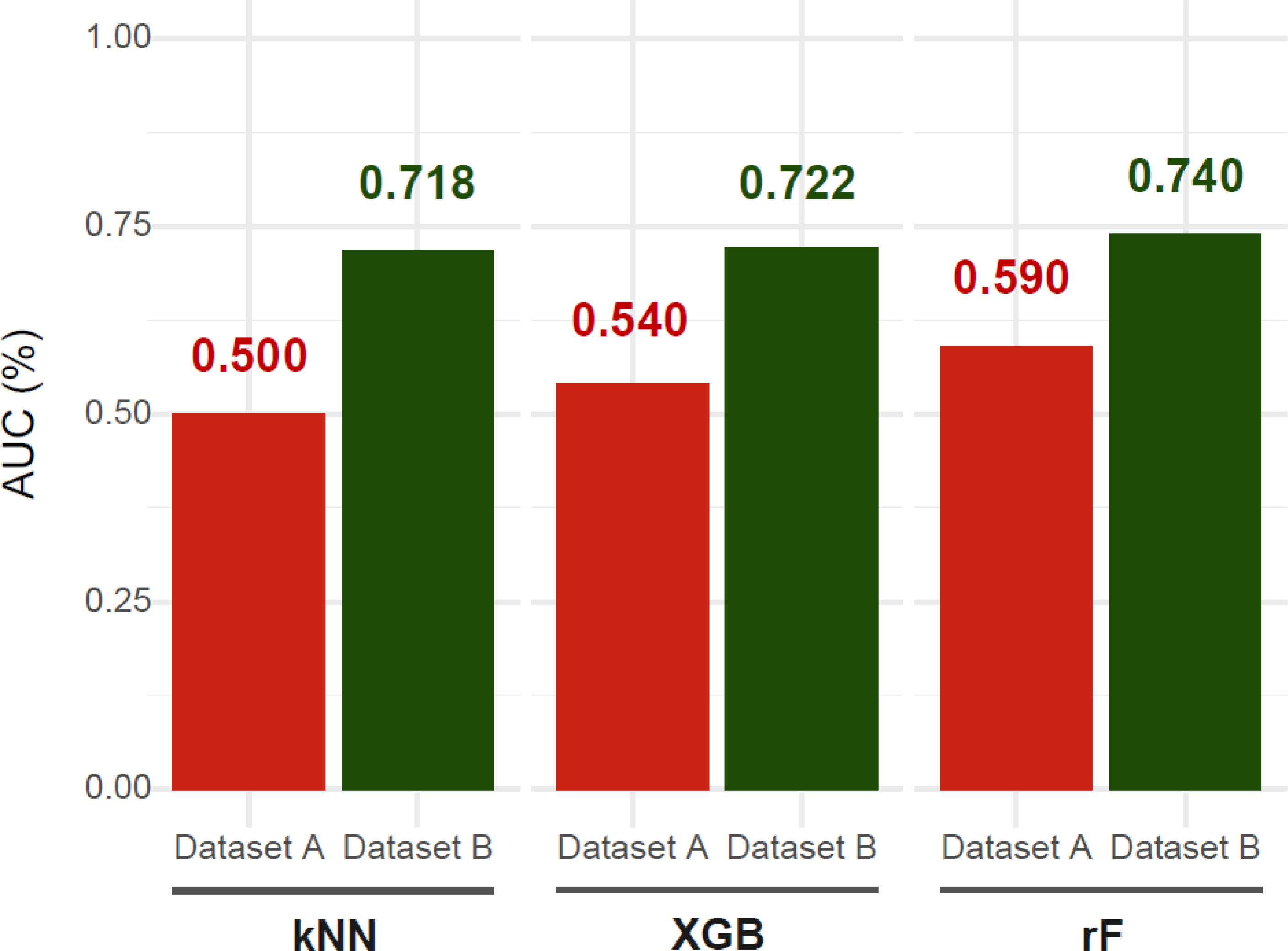
Results in terms of Area Under the Curve (AUC) between Dataset A and Dataset B in three Machine Learning algorithms (kNN, XGB, rF).

## DISCUSSION

Different morphokinetic patterns were detected by Machine Learning using aneuploid embryos as the contrary reference to Live-Birth embryos. As the algorithms, variables and Live-Birth embryos included in the ML analysis were the same for both Datasets, we think that lower results for Dataset “A” may be due to an increased Label Noise effect, suggesting that including aneuploid embryos could be more appropriate when building predictive models for embryo viability. We did not select euploid embryos since the limitations of blastomere biopsy on Day 3 would mean an increased risk of including mosaic embryos diagnosed as euploid. Non-concordance risk is inherent to the diagnosis technique as embryos comply with the Schrödinger cat paradox as, if the embryo was fully diagnosed there would not be the chance for transferring it, and if the whole embryo was not analysed, we cannot confirm the real genetic status of all of the non-biopsied cells. Moreover, euploidy does not necessarily mean an optimal embryo developmental potential. Similarly, the non-implanted group would assign a poor prognostic to potentially viable embryos due to factors unrelated to the embryo itself. Thus, determining the most suitable embryo categories for Artificial Intelligence algorithms is a matter of weights: despite mosaicism cannot be excluded from any category, we chose Live-Birth as it guarantees the desired embryo outcome and aneuploidy as an evidence of dysfunction within the embryo. We consider that this study should be performed again including trophectoderm biopsy instead of blastomere biopsy, and consequently, mosaic embryos could be taken into consideration by Machine Learning. It should be expected that the more the Label Noise is reduced, the better the accuracy of the predictive models. There is no clear evidence that morphokinetics could increase the cumulative success rate (Ahlstrom et al., 2016; Goodman et al., 2016, Apter et al., 2020). Higher inter- and intra- variability was described in embryo quality assessment by time-lapse monitoring, especially when considering features such as Pronuclear appearance and biomarkers after 8-cell stage (Sundvall et al., 2013; Martínez-Granados et al., 2017). For this reason, our model design consisted on using morphokinetic parameters until Day 3. Furthermore, our time-lapse & Machine Learning workflow contained 3 peculiar features which could play a role in improving TLT:

i. t0: time of ICSI was relativized for each oocyte, so dispersion in the first and last oocyte in a cohort was reduced to the minimum. It seems that the t0 used could increase the dispersion of t-times (Güell et al., 2019), although further studies should be performed to assess the impact of t0 in ML algorithms.
ii. Novel biomarkers: Nuclear fading at 2 cell-stage (tNf Cc2a, tNf Cc2b) and 4 cell-stage (tNf Cc3a) (Güell et al., 2018) could provide valuable data about the S and M phases (Aguilar et al., 2016) and make it possible to detect different phases length with a final overall cell cycle compensation which could be indistinguishable. These biomarkers are not included in nowadays Deep Learning image recognition although the expected variability between observers is expected to be low as these are abrupt events.
iii. Morphokinetic adjustments depending on morphodynamic phenomena: Irregular cleavages strongly correlate with impaired implantation (Milewski & Adjuk, 2017b). Annotating each MDp by type and cell cycle as a categorical feature could be suitable for estimating the risk and prognostic of the resulting embryo. With the current t-time annotation guidelines (Ciray et al., 2014) the annotation of t5 in tripolar mitosis at cell cycle 2 (from 2 to 5 cells), as an example, will cause a loss of all cell cycles traceability, even the hypothetic non-affected ones, unless particular adjustment should be applied. In fact, Herrero et al., 2013 recommends excluding MDp embryo from morphokinetic analysis. A schematic diagram of our guideline proposal and the consequences of traditional classic annotation of t5 in tripolar mitosis at cell cycle 2 can be displayed in Video 1.
iv. Cell lineage-based ML: ML variables included after feature extraction belonged to the A cell lineage (tNf Cc2a, t3, tNf Cc3a, t5). Future models could focus on B cell lineage and ensembled models could fit both cell lineage-based predictive models.

We are aware of the small dataset and we consider that it could be desirable to enlarge the sample size. Nevertheless, we assume that the minimum number of samples is beaten in this project since, as a rule of thumb, the sample size should be 10x the number of parameters in an algorithm (Nasir & Sassani, 2021). Moreover, a colossal embryo number would not necessarily mean better results as this could lead to a danger of increased disparity if the operators would follow each own protocol. According to this, automated parameter annotation (Khosravi et al., 2018) is becoming a promising solution as it would eliminate the human variability previously described. Furthermore, other non-invasive approaches, such as cell-free DNA analysis and metabolomics could be additionally performed for a more accurate embryo evaluation (Montag et al., 2013; Yang et al., 2014, Dominguez et al., 2015; Tejera et al., 2016).

This work was the first part of a global AI project, where we wanted to estimate the Implantation and Live-Birth Rate of transferred embryos predicted by an ensembled model of these algorithms. In conclusion, our ML results were improved when aneuploid embryos were taken into consideration. We believe that although prioritizing embryo transfer depending ML models could not lead to improved cumulative success rates, time-to-pregnancy could be reduced once the AI model was validated. Moreover, cell-cycle traceability could be advantageous for better explainability for patient counselling.

## ACKNOWLEDGMENT

We thank the internship IVF Lab students Laia, Estela, Óscar, Júlia, Jordina and Alejandra for their support during the data gathering and data cleaning process.

## RESEARCH DATA FOR THIS ARTICLE

Due to institutional requirement, data is unsuitable to post.

Data not available / The data that has been used is confidential

---

Table 1. Baseline characteristics of the study groups included in the datasets

## REFERENCES

1. Afnan, Michael Anis Mihdi and Rudin, Cynthia and Conitzer, Vincent and Savulescu, Julian and Mishra, Abhishek and Liu, Yanhe and Afnan, M. (2021). Ethical Implementation of Artificial Intelligence to Select Embryos in In Vitro Fertilization. Association for Computing Machinery. https://doi.org/10.1145/3461702.3462589

2. Aguilar, J., Rubio, I., Muñoz, E., Pellicer, A., & Meseguer, M. (2016). Study of nucleation status in the second cell cycle of human embryo and its impact on implantation rate. Fertility and Sterility, 106(2), 291–299.e2. https://doi.org/10.1016/j.fertnstert.2016.03.036

3. Ahlstrom, A., Park, H., Bergh, C., Selleskog, U., & Lundin, K. (2016). Conventional morphology performs better than morphokinetics for prediction of live birth after day 2 transfer. Reproductive BioMedicine Online, 33(1). https://doi.org/10.1016/j.rbmo.2016.03.008

4. Atwood, A. & Vadakkadath, S. (2016). The spatiotemporal hormonal orchestration of human folliculogenesis, early embryogenesis and blastocyst implantation. Molecular and Cellular Endocrinology, 430, 33–48. https://doi.org/10.1016/J.MCE.2016.03.039

5. Ciray, H. N., Campbell, A., Agerholm, I. E., Aguilar, J., Chamayou, S., Esbert, M., & Sayed, S. (2014). Proposed guidelines on the nomenclature and annotation of dynamic human embryo monitoring by a time-lapse user group. Human Reproduction, 29(12), 2650–2660. https://doi.org/10.1093/humrep/deu27

6. Curchoe, C., Flores-Saiffe Farias, A., Mendizabal-Ruiz, G., & Chavez-Badiola, A. (2020). Evaluating predictive models in reproductive medicine. Fertility and Sterility, 114(5), 921–926. https://doi.org/10.1016/J.FERTNSTERT.2020.09.159

7. Dimitriadis, I., Zaninovic, N., Badiola, A. C., & Bormann, C. L. (2022). Artificial intelligence in the embryology laboratory: a review. Reproductive BioMedicine Online, 0(0). https://doi.org/10.1016/J.RBMO.2021.11.003

8. Dominguez, F., Meseguer, M., Aparicio-Ruiz, B., Piqueras, P., Quiñonero, A., & Simón, C. (2015). New strategy for diagnosing embryo implantation potential by combining proteomics and time-lapse technologies. Fertility and Sterility, 104(4), 908–914. https://doi.org/10.1016/J.FERTNSTERT.2015.06.032

9. Fernandez, E., As, F., Mhm, C., Ds, C., Rcm, de S., Mfg, N., & Jc, R. (2020). Artificial intelligence in the IVF laboratory: overview through the application of different types of algorithms for the classification of reproductive data. Journal of Assisted Reproduction and Genetics, 37(10), 2359–2376. https://doi.org/10.1007/S10815-020-01881-9

10. Frénay, B., & Verleysen, M. (2014). Classification in the presence of label noise: A survey. IEEE Transactions on Neural Networks and Learning Systems, 25(5), 845–869. https://doi.org/10.1109/TNNLS.2013.2292894

11. Goodman, L. R., Goldberg, J., Falcone, T., Austin, C., & Desai, N. (2016). Does the addition of time-lapse morphokinetics in the selection of embryos for transfer improve pregnancy rates? A randomized controlled trial. Fertility and Sterility, 105, 275–285.e10. https://doi.org/10.1016/j.fertnstert.2015.10.013

12. Griffin, D., & Ogur, C. (2018). Chromosomal analysis in IVF: just how useful is it? Reproduction (Cambridge, England), 156(1), F29–F50. https://doi.org/10.1530/REP-17-0683

13. Güell, E., Ruiz, J., Ibarz, RM., Ibarz, JM. (2018). Morfocinética y sexo embrionario: aproximación a las implicaciones de la desaparición del núcleo blastomérico. MEDRE, Vol. 5 (Especial Congreso). 32° Congreso SEF (p. 91). Madrid, España.

14. Güell, E., Cura, J., Felip, L., Morono, E., González, O., Ibarz, J., Ibarz, R., Ruiz, J., & López, M. (2019). ¿T0 (hora de inicio time-lapse): puede modificar la valoración morfocinética de los embriones? In Mifsud, L., González, B., & Franco, Y. (Eds.), Revista ASEBIR: Vol. 24 N°2. X Congreso ASEBIR (pp. 91–92). Cáceres, España.

15. Herrero, J., Tejera, A., Albert, C., Vidal, C., De Los Santos, M. J., & Meseguer, M. (2013). A time to look back: Analysis of morphokinetic characteristics of human embryo development. Fertility and Sterility, 100(6). https://doi.org/10.1016/j.fertnstert.2013.08.033

16. Huang, B., Zheng, S., Ma, B., Yang, Y., Zhang, S., & Jin, L. (2022). Using deep learning to predict the outcome of live birth from more than 10,000 embryo data. BMC Pregnancy and Childbirth, 22(1), 1–7. https://doi.org/10.1186/S12884-021-04373-5/FIGURES/2

17. Irani, M., Reichman, D., Robles, A., Melnick, A., Davis, O., Zaninovic, N., Xu, K., & Rosenwaks, Z. (2017). Morphologic grading of euploid blastocysts influences implantation and ongoing pregnancy rates. Fertility and Sterility, 107(3), 664–670. https://doi.org/10.1016/J.FERTNSTERT.2016.11.012

18. Khosravi, P., Kazemi, E., Zhan, Q., Toschi, M., Malmsten, J. E., Hickman, C., Meseguer, M., Rosenwaks, Z., Elemento, O., Zaninovic, N., & Hajirasouliha, I. (2018). Robust Automated Assessment of Human Blastocyst Quality using Deep Learning. In bioRxiv. bioRxiv. https://doi.org/10.1101/394882

19. Kuhn, M., & Johnson, K. (2013). Applied predictive modeling. In Applied Predictive Modeling (Vol. 26). Springer. https://doi.org/10.1007/978-1-4614-6849-3

20. Martínez-Granados, L., Serrano, M., González-Utor, A., Ortíz, N., Badajoz, V., Olaya, E., Prados, N., Boada, M., Castilla, J. A., & Biology), on behalf of S. I. G. in Q. of A. (Spanish S. for the S. of R. (2017). Inter-laboratory agreement on embryo classification and clinical decision: Conventional morphological assessment vs. time lapse. PLOS ONE, 12(8), e0183328. https://doi.org/10.1371/JOURNAL.PONE.0183328

21. Milewski, R., Kuczyńska, A., Stankiewicz, B., & Kuczyński, W. (2017a). How much information about embryo implantation potential is included in morphokinetic data? A prediction model based on artificial neural networks and principal component analysis. Advances in Medical Sciences, 62(1), 202–206. https://doi.org/10.1016/j.advms.2017.02.001

22. Milewski, R., & Ajduk, A. (2017b). Time-lapse imaging of cleavage divisions in embryo quality assessment. In Reproduction (Vol. 154, Issue 2, pp. R37–R53). BioScientifica Ltd. https://doi.org/10.1530/REP-17-0004

23. Montag, M., Toth, B., & Strowitzki, T. (2013). New approaches to embryo selection. Reproductive BioMedicine Online, 27(5), 539–546. https://doi.org/10.1016/J.RBMO.2013.05.013

24. Morales, D., Bengoetxea, E., Larrañaga, P., García, M., Franco, Y., Fresnada, M., & Merino, M. (2008). Bayesian classification for the selection of in vitro human embryos using morphological and clinical data. Computer Methods and Programs in Biomedicine, 90(2), 104–116. https://doi.org/10.1016/J.CMPB.2007.11.018

25. Murugappan, G., Kim, J. G., Kort, J. D., Hanson, B. M., Neal, S. A., Tiegs, A. W.,

26. Nasir, V., & Sassani, F. (2021). A review on deep learning in machining and tool monitoring: methods, opportunities, and challenges. International Journal of Advanced Manufacturing Technology, 115(9–10), 2683–2709. https://doi.org/10.1007/S00170-021-07325-7

27. Osman, E. K., Scott, R. T., & Lathi, R. B. (2020). Prognostic value of blastocyst grade after frozen euploid embryo transfer in patients with recurrent pregnancy loss. F&S Reports, 1(2), 113. https://doi.org/10.1016/J.XFRE.2020.07.001

28. Popovic, M., Dheedene, A., Christodoulou, C., Taelman, J., Dhaenens, L., Van Nieuwerburgh, F., Deforce, D., Van den Abbeel, E., De Sutter, P., Menten, B., & Heindryckx, B. (2018). Chromosomal mosaicism in human blastocysts: the ultimate challenge of preimplantation genetic testing? Human Reproduction, 33(7), 1342–1354. https://doi.org/10.1093/HUMREP/DEY106

29. Riegler, M. A., Stensen, M. H., Witczak, O., Andersen, J. M., Hicks, S. A., Hammer, H. L., Delbarre, E., Halvorsen, P., Yazidi, A., Holst, N., & Haugen, T. B. (2021). Artificial intelligence in the fertility clinic: status, pitfalls and possibilities. Human Reproduction, 36(9), 2429–2442. https://doi.org/10.1093/HUMREP/DEAB168

30. Rosenwaks, Z. (2020). Artificial intelligence in reproductive medicine: a fleeting concept or the wave of the future? Fertility and Sterility, 114(5), 905–907. https://doi.org/10.1016/J.FERTNSTERT.2020.10.002

31. Rudin, C., Chen, C., Chen, Z., Huang, H., Semenova, L., & Zhong, C. (2021). Interpretable Machine Learning: Fundamental Principles and 10 Grand Challenges.

32. Simopoulou, M., Sfakianoudis, K., Maziotis, E., Antoniou, N., Rapani, A., Anifandis, G., Bakas, P., Bolaris, S., Pantou, A., Pantos, K., & Koutsilieris, M. (2018). Are computational applications the “crystal ball” in the IVF laboratory? The evolution from mathematics to artificial intelligence. Journal of Assisted Reproduction and Genetics, 35(9), 1545. https://doi.org/10.1007/S10815-018-1266-6

33. Swain, J., VerMilyea, M. T., Meseguer, M., Ezcurra, D., Ezcurra, D., Letterie, G., Sánchez, P., Trew, G., Nayot, D., Campbell, A., Huangv, I., Choma, J., Loewke, K., Piqueras, M. P., Nader, P., Schindler, M., Lippolis, E., Bohl, S., Kirsten, J., & Abshagen, D. (2020). AI in the treatment of fertility: key considerations. Journal of Assisted Reproduction and Genetics, 37(11), 2817–2824. https://doi.org/10.1007/s10815-020-01950-z

34. Tejera, A., Castelló, D., de los Santos, J. M., Pellicer, A., Remohí, J., & Meseguer, M. (2016). Combination of metabolism measurement and a time-lapse system provides an embryo selection method based on oxygen uptake and chronology of cytokinesis timing. Fertility and Sterility, 106(1). https://doi.org/10.1016/j.fertnstert.2016.03.019

35. Vermilyea, M., Hall, J. M. M., Diakiw, S. M., Johnston, A., Nguyen, T., Perugini, D., Miller, A., Picou, A., Murphy, A. P., & Perugini, M. (2020). Development of an artificial intelligence-based assessment model for prediction of embryo viability using static images captured by optical light microscopy during IVF. Human Reproduction, 1–15. https://doi.org/10.1093/humrep/deaa013

36. Victor, A. R., Griffin, D. K., Brake, A. J., Tyndall, J. C., Murphy, A. E., Lepkowsky, L. T., Lal, A., Zouves, C. G., Barnes, F. L., McCoy, R. C., & Viotti, M. (2019). Assessment of aneuploidy concordance between clinical trophectoderm biopsy and blastocyst. Human Reproduction, 34(1), 181–192. https://doi.org/10.1093/HUMREP/DEY327

37. Wang, R., Pan, W., Jin, L., Li, Y., Geng, Y., Gao, C., Chen, G., Wang, H., Ma, D., & Liao, S. (2019). Artificial intelligence in reproductive medicine. In Reproduction (Vol. 158, Issue 4). https://doi.org/10.1530/REP-18-0523

38. Weimar, C., ED, P. U., G, T., CJ, H., & NS, M. (2013). In-vitro model systems for the study of human embryo-endometrium interactions. Reproductive Biomedicine Online, 27(5), 461–476. https://doi.org/10.1016/J.RBMO.2013.08.002

39. Yang, Z., Zhang, J., Salem, SA., Liu, X., Kuang, Y. & Salem, RD. (2014). Selection of competent blastocysts for transfer by combining time-lapse monitoring and array CGH testing for patients undergoing preimplantation genetic screening: a prospective study with sibling oocytes. BMC Medical Genomics, 7(1). https://doi.org/10.1186/1755-8794-7-38

40. Zaninovic, N., & Rosenwaks, Z. (2020). Artificial intelligence in human in vitro fertilization and embryology. Fertility and Sterility, 114(5), 914–920. https://doi.org/10.1016/J.FERTNSTERT.2020.09.157

41. Zhan, Q., Sierra, E. T., Malmsten, J., Ye, Z., Rosenwaks, Z., & Zaninovic, N. (2020). Blastocyst score, a blastocyst quality ranking tool, is a predictor of blastocyst ploidy and implantation potential. F&S Reports, 1(2), 133–141. https://doi.org/10.1016/J.XFRE.2020.05.004

42. Zhu, X., & Wu, X. (2004). Class Noise vs. Attribute Noise: A Quantitative Study. Artificial Intelligence Review 2004 22:3, 22(3), 177–210. https://doi.org/10.1007/S10462-004-0751-8

